# Flexible predictive processing during face perception

**DOI:** 10.64898/2026.07.29.741445

**Authors:** Francisco Gutiérrez-Blanco, Sergio Gaspar, Ana F. Palenciano, Carlos González-García, María Ruz

**Author notes:** Correspondence should be addressed to: María Ruz, Mind, Brain and Behavior Research Center (CIMCYC), Department of Experimental Psychology, University of Granada, Granada, Spain.

## Abstract

Perceptual expectations facilitate perception of expected information by shaping neural coding. However, how the cognitive demands of different tasks may modulate this effect remains unclear. Here, we investigated whether and how expectations derived from sex/gender stereotypes impact behavioral and neural face processing as a function of task goals. Fifty-one participants categorized either the emotional expression (happy vs. angry) or the sex (male vs. female) of faces while mouse trajectories and electroencephalography (EEG) were recorded in separate sessions. Expectations were manipulated through the congruency between facial sex and emotional expression (congruent: angry male, happy female; incongruent: happy male, angry female). Behavioral analyses revealed that incongruent faces elicited larger trajectory deviations, longer movement dynamics and slower reaction times than congruent faces. Representational Similarity Analyses (RSA) showed that face congruency modulated coding geometries of both mouse trajectories and EEG, whereas multivariate classifiers revealed progressive impact of congruency from early stages of face neural coding. Importantly, all measurements revealed stronger congruency effects during emotion than during sex judgments, indicating that the influence of expectations is flexibly tailored to task demands. Furthermore, congruent faces exhibited enhanced decoding performance and reached peak neural discriminability earlier than incongruent ones. Finally, EEG–mouse fusion analyses revealed shared representational dynamics primarily explained by task demands. Together, these findings indicate that expectations facilitate face processing in a task-dependent manner and suggest that predictive facilitation operates, rather than by increasing the rate of evidence accumulation, by accelerating the emergence of neural coding patterns.

**Significance Statement:** Perception is shaped by the combination of sensory input and prior expectations, including socially acquired sex/gender face stereotypes. Learning how these expectations influence perception and how they are flexibly adjusted to current behavioral goals is essential for predictive coding models. Combining EEG and mouse tracking, we show that emotion judgments are more susceptible to expectations than sex categorization, and that expectations accelerate the emergence of neural coding patterns rather than simply increasing the rate of evidence accumulation. These findings characterize the neural mechanism through which predictive processing adapts to task demands and demonstrate that the influence of social stereotypes on face perception is dynamically regulated rather than automatically imposed, providing new insights into when social biases are most likely to shape human perception.

## 1. Introduction

Predictive processing models (Friston, 2005; Summerfield & Egner, 2009) propose that perceptual expectations are conveyed through top-down feedback from higher cortical regions, reshaping the neural representational space of lower-level cortical regions to facilitate the encoding of expected information (Auksztulewicz & Friston, 2016; Egner et al., 2010; Kok et al., 2017; Nigam & Schwiedrzik, 2024). Being face perception a prominent model for studying predictive processing, expectation effects have been reported across identity (Robinson et al., 2020), ethnicity (Hehman et al., 2014), emotional expression (Baseler et al., 2014) or sex/gender stereotypes (Barnett et al., 2021) of faces. Importantly, these processes usually unfold under different goals, which impose distinct cognitive demands. Investigating how expectations interact with tasks goals is essential to explaining how the brain flexibly adapts perception to changing behavioral demands. In this work, we studied whether and how demands to judge either the emotion or the sex of faces modulate the effect of expectations based on sex/gender stereotypes during dynamic face perception, using a combined mouse tracking and electroencephalography (EEG) approach.

Building on stereotypical associations linking male faces to anger and female faces to happiness (Hess et al., 2000, 2004, 2022), multiple studies have investigated the influence of stereotypes on face perception (Barnett et al., 2021; Brooks et al., 2018; Stolier & Freeman, 2017). At the behavioral level, stereotype-congruent faces (angry male and happy female) are categorized faster and more accurately than incongruent ones (happy male and angry female; Becker et al., 2007). In addition, continuous behavioral measures such as mouse tracking capture the magnitude and temporal dynamics of these expectations, with larger Maximum Absolute Deviations (MAD) and longer MAD-time windows for incongruent than for congruent faces (Barnett et al., 2021; Freeman & Ambady, 2010; Gutiérrez-Blanco et al., 2025). Moreover, Representational Similarity Analysis (RSA) shows that these effects are reflected on the spatiotemporal patterns of the mouse movements at intermediate stages of face processing (Gutiérrez-Blanco et al., 2025).

Neuroimaging studies have consistently associated incongruent faces with increased neural activity (Stolier & Freeman, 2017), and more recently, enhanced top-down functional connectivity between higher-order orbitofrontal regions involved in predictive processing and lower-level face-sensitive fusiform cortex (Barnett et al., 2021). However, the limited temporal resolution of functional magnetic resonance imaging (fMRI) prevents direct exploration of the temporal dynamics of sex/gender expectations during face perception. Although, to our knowledge, no EEG study has directly addressed this question, related studies employing univariate approaches have reported inconsistent findings, with expectation effects emerging either during early perceptual stages indexed by the N170 potential (Freeman et al., 2010) or during later perceptual and decision-related stages indexed by the P300 (den Ouden et al., 2023; Feuerriegel et al., 2018; Summerfield et al., 2011). Multivariate classifiers provide a powerful framework to characterize the temporal dynamics of predictive expectations and determine whether expectation-driven facilitation arises through earlier neural decoding, faster evidence accumulation, or both.

Importantly, expectations unfold under different task goals that engage distinct cognitive processes. Consistent with this, facial sex and emotional expression show asymmetric interference during categorization: emotion judgments are influenced by facial sex, whereas sex judgments are comparatively resistant to emotional expression (Atkinson et al., 2005; Karnadewi & Lipp, 2011). More recently, Gandolfo et al. (2025) showed that sex and emotion rely on partially distinct processing mechanisms, with emotion showing greater dependence on top-down influences than sex processing. This is consistent with influential models proposing partially dissociable pathways for processing invariant, sex, and variable, emotion, facial information (Bernstein & Yovel, 2015; Haxby et al., 2000; Robinson et al., 2020). Together, these findings suggest that task goals may differentially shape the influence of sex/gender stereotype expectations during face perception.

We combined EEG and mouse tracking to investigate how task goals shape the temporal dynamics of sex/gender stereotype expectations during face perception. Based on previous behavioral and neuroimaging findings, we predicted stronger expectation effects during emotion than sex categorization, reflecting the greater contribution of top-down processing to emotional processing. We further tested whether expectations facilitate face processing by advancing the emergence of neural decoding, increasing the rate of evidence accumulation, or both. Finally, we examined whether expectation effects are expressed through convergent representational dynamics across EEG and mouse trajectories.

## 2. Methods

### 2.1. Participants

Fifty-one volunteers were recruited from the University of Granada (30 women, 21 men; mean age = 22.37; range = 18 - 30). They all signed a consent form approved by the local Ethics Committee for Human Research (ref 1584/CEIH/2020) and received compensation of 30 Euros. The sample size was determined based on previous studies that used a similar methodology to investigate the effect of expectations on face processing (Barnett et al., 2021; Stolier & Freeman, 2016, 2017). All participants were included in the analyses of the mouse trajectories while five were excluded from the EEG analysis due to excessive signal noise (with more than 30% discarded trials), with a final EEG sample of 46 participants.

### 2.2. Apparatus and stimuli

The experiment was conducted on a Windows 10 computer with LDC display (1920 x 1080-pixel resolution and 60 Hz refresh rate), situated at 60 cm from the participants.

A total of 60 photographs of Caucasian faces were extracted from the Chicago Face Database (Ma et al., 2015), corresponding to 30 different identities (15 male and 15 female), each expressing either anger or happiness. For the entire sample of participants, twenty face identities were randomly selected for the experimental sessions (from a total of 40 faces), while the remaining were used for practice. Faces were transformed to grayscale, and their luminance and contrast levels were equated using the luminance and contrast values of a randomly chosen face as a criterion, again for the whole set. The central part of the faces was extracted using an oval mask and their size and position were adjusted to three reference points on the mask, two for the eyes and one for the nose. This process was carried out with Adobe Photoshop software. The combination of sex/gender and emotion created the categories of congruent (angry male and happy female) and incongruent (happy male and angry female) faces, in accordance with expectations based on sex/gender stereotypes (Barnett et al., 2021; Gutiérrez-Blanco et al., 2025; Hess et al., 2000, 2004).

### 2.3. Experimental design and procedures

Participants were required to complete two experimental sessions, with an interval of 1-3 days between them. The first employed a mouse tracking paradigm, while the second entailed electroencephalography (EEG) recording. Both were performed at the Mind Brain and Behavior Research Centre (CIMCYC) of the University of Granada.

The mouse tracking task was programmed using OpenSesame, with the Mousetrap plugin (Kieslich et al., 2017). The screen background and text elements (situated within the response button) were displayed in black, while the rest (response buttons and fixation cross) appeared in grey. The EEG task was programmed in MATLAB (r2023) with Psychtoolbox (Brainard, 1997). The background of the screen was grey to reduce the contrast with the faces, while the text elements were displayed in white.

In both sessions (mouse tracking and EEG), participants were instructed to categorize, in different blocks, either the sex (male or female) or the emotion (angry or happy) of faces that were congruent (angry male and happy female) or incongruent (happy male and angry female) with expectations based on sex/gender stereotypes. In the mouse task, participants had to respond by clicking on the response buttons located at the two upper corners of the screen. The EEG task was adapted to minimize eye movements, in this case, participants had to use the computer keys to respond. Prior to the experimental sessions, participants were instructed on the sequence of events and completed two practice blocks, one of sex and one of emotion judgments. Practice of a localizer block (see below) was also included in the EEG task. In each practice block, 20 faces were randomly presented with a feedback signal at the end of each trial indicating whether the response was correct or incorrect. Responses and block order were counterbalanced across participants.

#### Mouse task

At the beginning of each trial, a fixation cross appeared for 500 ms at the center of the screen (degrees of visual angle, dva 0.9°, 0.9°), followed by a screen displaying a START button at the bottom center of the screen (dva; 16.7°, 10.3°) and two response buttons at the left and right top corners from the START button (dva; 16.7°, 10.3°). These contained the labels ANGRY - HAPPY, in the emotion task and MALE - FEMALE, in the sex task. Once the participants pressed the START button, it disappeared and, 200 ms afterwards, the response screen displayed a photograph (dva; 7.8°, 11.4°) at the center of the screen, together with the response buttons. The trial concluded when participants clicked on one of the response buttons (see Figure 1B).

**Figure 1.**
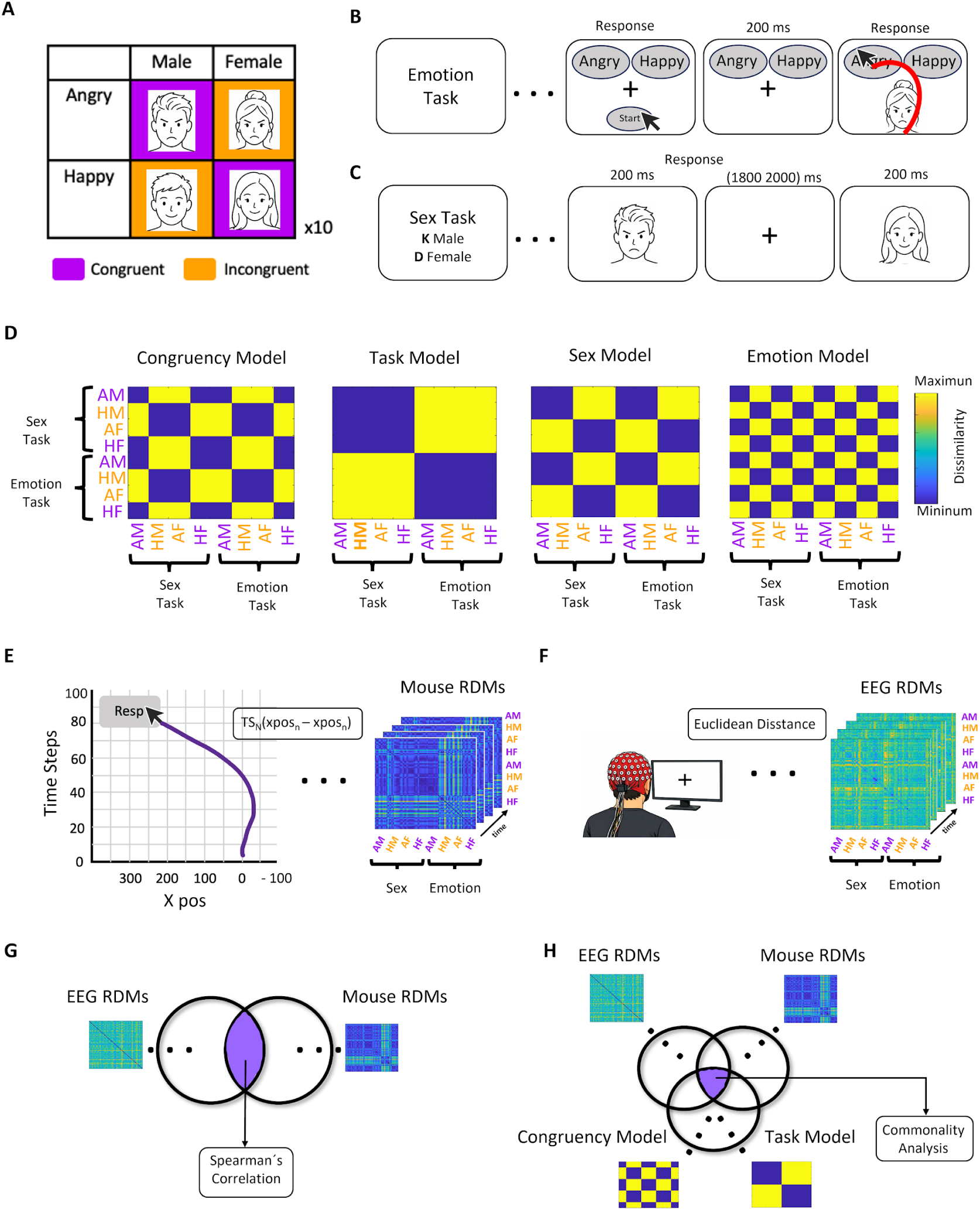
Experimental design and representational similarity analysis pipeline. **(A)** Stimulus space illustrating face congruency according to sex/gender stereotypes. Faces combined facial sex (male, female) and emotional expression (angry, happy), generating congruent stimuli (angry male, happy female) and incongruent stimuli (happy male, angry female). **(B)** Mouse-tracking task sequence. Participants performed two task types in separate blocks: an emotion categorization task (angry vs. happy) and a sex categorization task (male vs. female). Responses were given by moving the cursor from a start position toward one of two response alternatives located at the upper corners of the screen. **(C)** EEG task sequence. Participants performed the same two task types in separate blocks (emotion categorization and sex categorization), responding by pressing one of two response keys following face presentation. **(D)** Theoretical representational dissimilarity matrices (RDMs) used in model-based representational similarity analysis (RSA). RDMs were designed to reflect the expected representational distances (dissimilarities) between face stimuli as a function of Congruency (congruent vs. incongruent), Task type (emotion vs. sex categorization), **Sex** (male vs. female), and Emotion (angry vs. happy). Distances were coded as low (0) or high (1) dissimilarity. **(E)** Mouse-trajectory RSA pipeline. Temporally normalized trajectories were represented as time-resolved horizontal cursor positions (x-axis). Empirical RDMs were computed at each time point from pairwise differences between trajectory patterns across experimental conditions. **(F)** EEG RSA pipeline. Time-resolved empirical neural RDMs were generated by computing Euclidean distances between multivariate EEG activity patterns across electrodes for all experimental conditions. **(G)** EEG–mouse representational fusion. Similarity between mouse and EEG representational geometries was estimated using Spearman correlations between vectorized RDMs across all combinations of mouse and EEG time points, yielding a temporal generalization matrix of representational similarity. **(H)** Model-based commonality analysis. Shared variance between EEG and mouse representational geometries was decomposed to estimate the variance uniquely explained by theoretical models of Congruency and Task type, quantifying their contribution to the representational overlap between neural activity and mouse trajectories.

#### EEG task

The sequence of trial events was the following (see Figure 1B). First a fixation cross was displayed at the center of the screen for a variable time interval between 1800 and 2000 ms (dva; 0.9°, 0.9°). Then a face appeared at the center of the screen (dva; 7.8°, 11.4°) for 200 ms and participants had a maximum of 800 additional milliseconds to judge, depending on the task block, its sex (male or female) or emotion (happy or angry) by pressing the corresponding keys (D or K).

In each session participants performed a total of 32 blocks (16 of each task). In each block, all faces were presented in random order, resulting in 40 trials per block and a total of 1280 trials per session. The order of presentation of the blocks was designed to ensure that there were the same number of contiguous same and different blocks. Our design also balanced the position of the response options across blocks, such that each type of task had the same number of blocks in each disposition (e.g., female-left, male-right in mouse task and female-D, male-K in EEG task). The duration of the mouse task was 50-60 minutes, while the time to complete the EEG task was between 80 to 95 minutes.

Both tasks (Mouse and EEG) had a 2×2 within-participant experimental design, with Face Congruency (congruent vs. incongruent) and Task type (sex vs. emotion categorization) as main factors.

Additionally, a third type of block (i.e., localizer blocks) was collected during the EEG task to obtain data during face processing isolated from the influence of task demands associated with the experimental blocks. Here participants responded by pressing the "B" key only when a face appeared rotated 180°. The sequence of events was identical to that designed for the other two types of blocks. Participants performed a total of 16 localizer blocks, in which all faces were randomly presented plus 4 rotated faces (one of each of the four types of stimuli, chosen randomly). These data, however, were not employed for the present analyses and will not be discussed further.

### 2.4. Data Analyses

#### 2.4.1. Mouse Trajectories Acquisition and Preprocessing

Trajectories were recorded at a sampling rate of 5 ms. The coordinates of the mouse at the vertical and horizontal axes of the screen were collected from the onset of the movement until the response.

Preprocessing of the trajectories was conducted in RStudio (R Core Team, 2016) with the R package mousetrap (Kieslich et al., 2019), following the protocol proposed by Wulff et al. (2026). Only trials with correct responses (average 98.4%) and faster than 2000 ms (98.9%) were employed. All trajectories were realigned to the same left response position and were matched to the same start positions at the [0,0] coordinate axes. To identify and exclude outliers, the following procedure was employed: (1) trajectories were normalized longitudinally so that they were represented by the same number (100) of spatially equidistant points; (2) the normalized trajectories were clustered based on their similarity (Euclidean distance) to five different prototypes previously designed by Wulff et al (2026); and (3) trials with trajectories that exhibited a standard deviation greater than or equal to 2 to their prototype were excluded (4.72%). Following pre-processing, the maximum absolute deviation (MAD) points and the time intervals between the onset of the movement and the MAD point (MAD time) were extracted for each trial.

#### 2.4.2. EEG data Acquisition and Preprocessing

Behavioral data (accuracy and reaction times, RTs) were recorded during the EEG task. Prior to analysis, trials with incorrect responses and RT outliers (±2.5 SD from each participant’s mean RT) were excluded (average 8.63%; range 2.58-23%).

Brain electrical activity was recorded with high-density EEG of 64 electrodes (BrainVision actiCap Slim) sampled at 1000 Hz and referenced to FCz. Electrodes TP10 and TP21 were positioned on the inferior and lateral areas of the right eye of the participants to track eye blinks and movements. To provide optimal recording conditions, electrode impedances were maintained below 10 kΩ.

EEG data were preprocessed in MATLAB (r2023) with the EEGLAB toolbox (Delorme & Makeig, 2004), using custom MATLAB scripts (https://github.com/CIMCYC/EEG/tree/main/MATLAB/PREPROCESSING/EEGLAB/L%C3%93PEZ%2C%20D). The EEG signal was resampled at 256 Hz and filtered above 0.1 Hz and below 120 Hz, also with notch bandpass at 50 and 100 Hz. Noisy channels detected during recording were removed, and epochs locked to face onset, from -200 to 1000 ms, were extracted from all trials. Artifacts derived from blinks and eye movements were detected through visual inspection and the ICLabel tool (Pion-Tonachini et al., 2019) after the Independent Component Analysis (ICA), resulting in the removal of 1.38 components on average (range 0-3) per participant. This was followed by an automatic trial rejection procedure based on the following factors: 1) abnormal spectrum, removing trials where the spectrum differed significantly from the baseline in +- 50 dB in the 0-2 Hz frequency window and -100 dB or +25 dB in the 20-40 Hz frequency band, artifacts previously associated with eye activity and muscle movements, respectively; 2) improbable data, removing trials with voltage values +-6 SD from the mean probability distribution; and 3) extreme amplitude values, removing trials with ±150 μV on at least one electrode (see Lopez-García, 2022; Pena et al., 2025; Peñalver et al., 2023 for similar preprocessing protocols). The discarded channels were interpolated (average 0.17, range 0-2), and the activity was re-referenced to a common average. Then, trials were corrected to the baseline set between -200 to 0 ms and those with incorrect responses were discarded. After preprocessing, the average of observations per participant was 85% (range 70-95%).

#### 2.4.3. Behavioral Analysis

##### Mixed Effects Models

To investigate whether the degree of curvature (MAD) and the time interval between the start of the movement and the MAD point (MAD time) in the mouse trajectories and behavioral performance (RT) in the EEG session were modulated by Congruency and Task Type, three independent mixed effect models were implemented, one for each dependent variable. These were performed in Rstudio with the *lmer* library. Post-hoc analyses of interaction effects were implemented with the *marginaleffects* library, applying the Bonferroni method to address multiple comparisons. Prior to the main analyses, and with the aim of optimizing the efficiency and fit of the mixed models to the 2×2 experimental design, a dichotomous coding system was implemented (see DeBruine and Barr, 2021), with values of -0.5 and 0.5 for the two levels of each variable of interest (Congruency: congruent = -0.5 and incongruent = 0.5; Task type: sex = -0.5 and emotion = 0.5).

To determine the design and structure of the mixed models, an identical procedure was followed in each of the analyses (see Supplementary Material). First, the levels of the variables Congruency (congruent vs. incongruent), Task type (sex vs. emotion) and their interaction were set as fixed effects. The random effects’ structure was designed by testing the goodness of fit between two models: one that considered the stimuli and participants as random factors and one that considered the stimuli, participants and the interaction between them. A likelihood ratio test revealed that the latter model presented a significantly better fit (all *ps* < .001). The final models were designed by testing all possible combinations of random slopes (combination between fixed and random effects) and selecting the best-fitting model based on the lowest AIC value (see following Formulas).

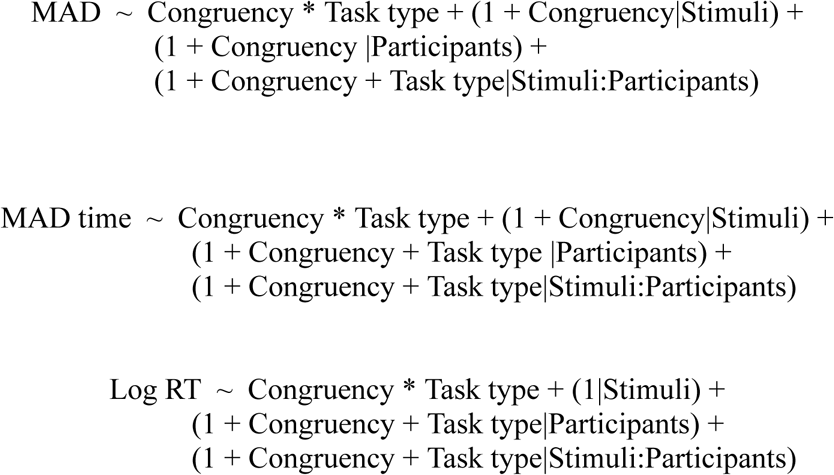

#### 2.4.4. Mouse and EEG data Analysis

##### Model-Based Representational Similarity Analysis (RSA)

We applied a time-resolved model-based RSA to characterize the representational geometry underlying spatiotemporal mouse trajectory patterns (Gutiérrez-Blanco et al., 2025) and brain electrical activity (Pena et al., 2025; Peñalver et al 2023; Kriegeskorte et al., 2008) as a function of our variables of interest. RSA quantifies representational structure by comparing empirical and theoretical representational dissimilarity matrices (RDMs). Empirical RDMs capture pairwise dissimilarities across experimental conditions, whereas theoretical RDMs encode the dissimilarity structure predicted by specific hypotheses about information representation. Analyses were implemented in MVPAlab (López-García et al., 2022; https://github.com/dlopezg/mvpalab).

Four theoretical RDMs were designed to reflect the expected distances between all individual face stimuli based on Congruency (congruent and incongruent), Task type (sex and emotion), Sex (male and female) and Emotion (angry and happy; see Figure 1D). This design included all face items in both tasks, with symmetrical 80×80 matrices (corresponding to the 40 different faces across the two task settings). The distances between conditions were coded as 0 (minimum dissimilarity) or 1 (maximum dissimilarity).

Mouse empirical RDMs were extracted as follows. Trajectories were temporally normalized so that they spanned 103 temporally equidistant time steps. For each time step, the mean horizontal cursor position (xpos; measured in screen pixels) was calculated across trials for each of the 80 experimental conditions. RDMs were then computed by calculating the pairwise differences in xpos between all experimental conditions, resulting in RDMs of 80×80 for each temporal step (Gutiérrez-Blanco et al., 2025; Koenig-Robert et al., 2024). This process was repeated for each individual participant (see Figure 1E).

EEG empirical RDMs were computed as follows. For each participant, neural activity patterns (raw voltage amplitude, across electrodes) per face condition were trial-averaged and centered around 0 by subtracting, for each electrode, the mean response across experimental conditions (Pena er al., 2025). And at each time point within the epoch (-200 to 1000 ms), an 80×80 RDM was constructed by calculating the Euclidean distance between averaged activity patterns from all possible pairs of face conditions.

Two independent, although identical model-based RSAs were implemented, one for mouse trajectories and one for EEG data. First, the empirical and theoretical RDMs were vectorized, removing the diagonal and the upper triangle values. This process avoids inflated correlation effects between matrices and improves the performance of the statistical analyses (Ritchie, Bracci & Beeck, 2017; Peñalver et al., 2023, 2024). Then, to estimate the proportion of variance in the empirical RDMs explained by the different theoretical models, multiple linear regressions were run at each time point. The 4 theoretical RDMs (also vectorized after removing diagonal and upper triangle) were considered as regressors, while the empirical RDMs were the dependent variables. These analyses returned unique beta weights and the corresponding t-values for each model, time point and participant.

Finally, we performed a non-parametric cluster-based permutation method to infer statistical significance at the group level. First, we conducted 100 permutations per individual participant. For each of these, the structure of the theoretical RDMs was randomized, after removing the diagonal/lower triangle, and included as regressors in the multiple linear regression models. We then generated 10^5^ permutations at the group level. In each of these, the result of one permutation per participant was randomly selected and averaged across them. This process generated null distributions of group-level t-values centered at around 0, which were used to estimate the above and below thresholds of the empirical probability distribution for each time point and model, considering a two-tailed 95% percentile (97.5th in each tail). To determine the minimum cluster size for statistical significance, we grouped continuous and significant time points in each of the permutations and selected those with a size above the 95th percentile. Finally, clusters of contiguous suprathreshold time points were then identified, and their statistical significance was assessed using a cluster-level false discovery rate (FDR) correction (α = 0.05), as described in Stelzer et al. (2013) and López-García et al. (2022).

To determine whether the activity patterns reflecting the congruency of faces with sex/gender stereotypes was modulated by task demands, we followed the procedure described below, separately for each of the data modalities. First, we conducted two additional time-resolved model-based RSAs, one for each task type. Empirical RDMs (mouse trajectories and EEG) were computed following the same procedures described above, except that the analyses were restricted to the 40 face stimuli presented within each task type, resulting in RDMs of 40×40. The time-resolved model-based RSAs were run including a single regressor of a 40×40 theoretical RDM reflecting the congruence of faces with sex/gender stereotypes. Group-level statistical significance was assessed using the same non-parametric cluster-based permutation approach. Second, we calculated the differences between the resulting t-values (emotion RSA - Sex/Gender RSA) at each time point and participant individually. To establish statistical significance at the group level, we performed a non-parametric cluster-based permutation. In this case, we retrieved the 100 permutations generated in the two previous RSAs (emotion and sex tasks) and calculated their differences at each time point and for each participant. This allowed us to calculate 10^5^ permutations at the group level, generating a null distribution of differences centered around 0 and estimating the above and below thresholds of the empirical probability distribution for each time point, again considering a two-tailed 95% percentile. The rest of the procedure was identical to that explained above.

##### Mouse-EEG Fusion

To explore the similarity between the geometric representational structure of the spatiotemporal patterns of mouse trajectories and EEG data, we performed a time resolved simple fusion analysis of mouse and EEG data, following and adapting previous work by Hebart et al. (2018), Peñalver et al. (2024) and Koenig-Robert et al. (2024).

We retrieved the vectorized RDMs derived from the previous mouse and EEG RSAs and selected only those participants included in the EEG analyses (n = 46). A moving average filter was applied every 5 time points to the EEG RDM. For every time bin, t_n_, the features of the previous and the following data points were averaged, so that t_n_ = (t_n-2_ + t_n-1_+ t_n_+ t_n+1_+ t_n+2_)/5. Then we computed Spearman correlations between vectorized mouse and EEG RDMs for every possible combination of time points. This resulted in a temporal generalization matrix of 102 (mouse-time) × 307 (EEG-time, see Figure 1G). This approach allowed estimating when the representational structures underlying both modalities exhibited similarity (Koenig-Robert et al., 2024). This procedure was repeated separately for each participant.

At the group level, statistical inference was performed using a non-parametric cluster-based permutation approach, as described in Koenig-Robert et al. (2024). Because correlation coefficients are bounded and exhibit non-normal sampling distributions, individual temporal generalization matrices of correlations were Fisher z-transformed prior to statistical testing. This transformation stabilizes variance and improves the normality of the sampling distribution at the group level. For each pair of possible combinations, a one-sample t-test against zero was performed across participants, yielding a two-dimensional map of t-values and associated p-values. Clusters were defined by applying a threshold of p < .01 (uncorrected) to the resulting p-value map in the two-dimensional space. Statistical significance was evaluated using a sign-flip permutation procedure (10,000 permutations). Under the null hypothesis of zero mean effect, the sign of each participant’s Fisher-transformed matrix was randomly inverted, and the entire statistical procedure (t-test and cluster identification) was recomputed. For each permutation, the maximum cluster mass (defined as the sum of t-values within each cluster) was retained to build a null distribution of the largest cluster statistic expected by chance. Observed clusters were considered significant if their cluster statistic exceeded the 95th percentile of this null distribution, thereby controlling the FWER at α = .05.

To determine which portion of the shared variance between the representational geometric structures of mouse and EEG patterns was explained through time by Congruency and Task, we employed a time-resolved model-based commonality analysis (Koenig-Robert et al., 2024; see Figure 2D). First, we applied the same preprocessing pipeline described in the time-resolved simple fusion analysis, including, in this case, the vectorization of the theoretical Congruency and Task type RDMs. Notably, unlike the previous analyses, the present commonality analysis was restricted to the Congruency and Task models, as these corresponded to the factors of primary theoretical interest and reduced the computational burden. Second, the commonality coefficient (C) was computed as the difference between (i) the squared semi-partial Spearman correlation between EEG and mouse RDMs while controlling for all theoretical RDMs except the model of interest (R²ₐ), and (ii) the squared semi-partial Spearman correlation between EEG and mouse RDMs while controlling for all theoretical RDMs simultaneously (R²ᵦ). The analysis was performed for every combination of mouse-time and EEG-time points, resulting in a 102 × 307 time–time commonality matrix per model. We repeated the process for each model and participant.

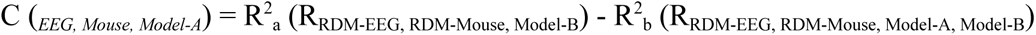

**Figure 2:**
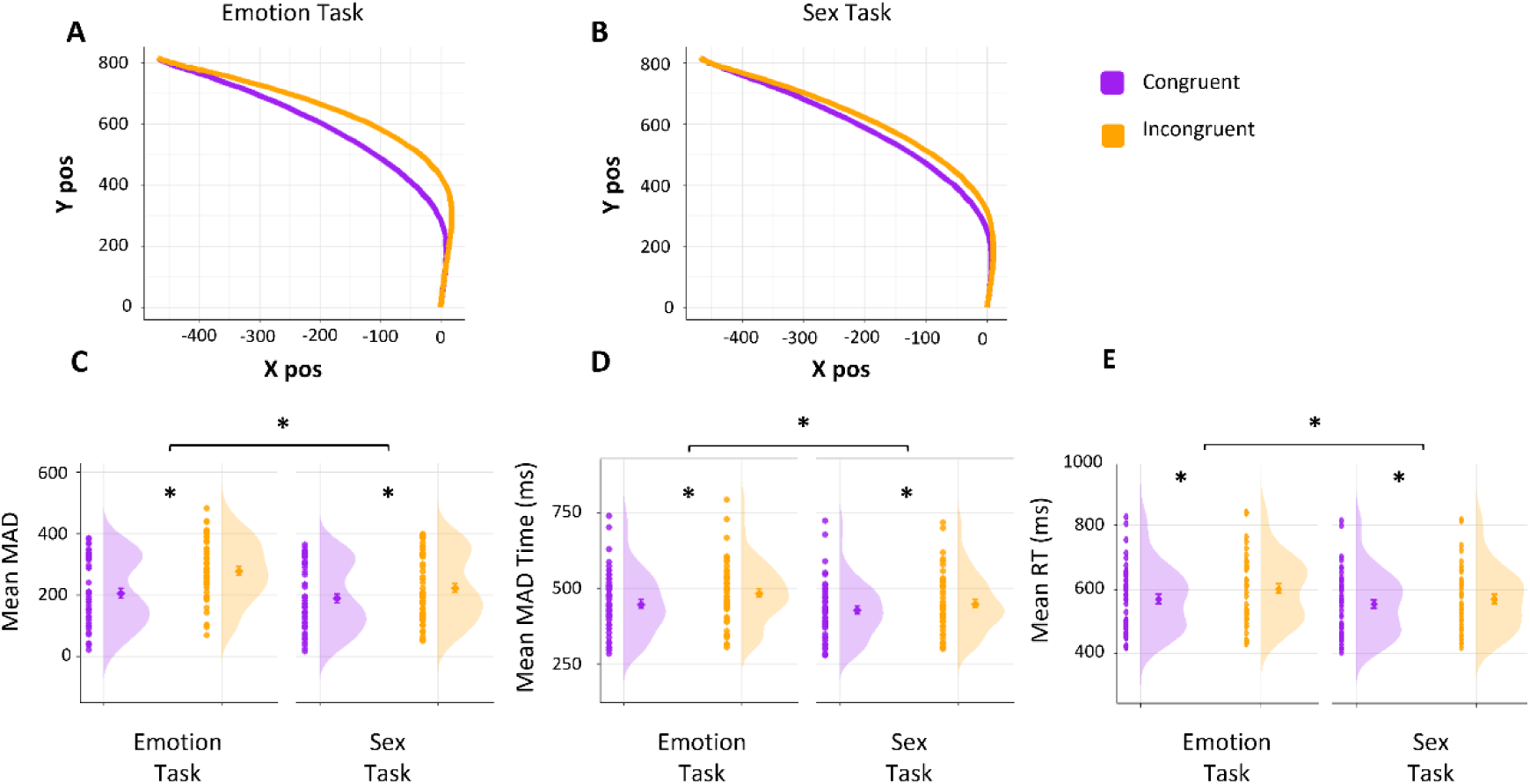
Behavioral results. **(A–B)** Average mouse trajectories (xpos and ypos, in screen pixels) during emotion categorization (**A**) and sex categorization (**B**) for congruent and incongruent faces. **(C–D)** Mean maximum absolute deviation (MAD; **C**) and MAD-time (**D**) as a function of Task type and Congruency. **(E)** Mean reaction times (RTs) recorded during the EEG task across task and congruency conditions. Individual points represent participant-level values, central markers indicate condition means, error bars represent ± SEM, and violin plots depict the distribution of participant-level values. Asterisks indicate statistically significant differences (*p* < .001).

Group-level inference was conducted using the same non-parametric cluster-based permutation procedure described for the simple fusion analysis. However, in this case no Fisher z-transformation was applied.

##### Time resolved Multivariate Classifiers

To examine when in time the neural activity patterns encode the congruency of faces with expectations based on sex/gender stereotypes, Task type and the Sex and Emotion of the faces, we carried out time-resolved Multivariate Pattern Analyses (MVPA) (Grootswagers et al., 2018) employing the MVPAlab Toolbox (López-García et al., 2022; https://github.com/dlopezg/mvpalab). While RSAs allow estimating the neural representation space of specific information through the comparison of RDMs and theoretical models, MVPA tests whether multivariate neural activity patterns contain sufficient information to discriminate between experimental conditions in the time domain (Grootswagers et al., 2018; Kriegeskorte et al., 2008; Pena et al., 2025; Peñalver et al., 2023).

We trained and tested classification algorithms to discriminate the EEG activity patterns (raw voltage values across channels, per each time point) of trials from the two levels of each of our independent variables (Congruency: congruent vs incongruent faces, Task type: sex vs emotion categorization, Sex: male vs female and Emotional expression: angry vs happy). Prior to the analysis, to increase the signal-to-noise ratio and to reduce computational costs, the epochs for each trial were sub-sampled with a step size of 3 time points (Pena et al., 2025; Peñalver et al., 2023). The whole set of trials per condition and participant was divided into five equal parts to ensure that the subsequent cross-validation procedure was balanced. Test and training data were then normalized across each of the folds (King and Dehaene, 2014), as follows:

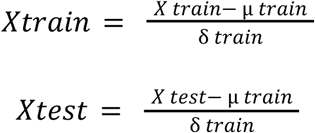

In the formula μ train and σ train indicate the mean and standard deviation of the training set. This process adjusts the range of the raw EEG data to a common scale without distorting the differences in the ranges of values. Afterwards, data were smoothed using a moving average filter with a length of 3 time points (Grootswagers et al., 2017).

Within this 5-fold cross-validation procedure, we implemented a Linear Discriminant Analysis (LDA; Grootswagers et al., 2017; Pena et al., 2025). We employed the Area Under the Curve (AUC) as a measure of classifier performance, as it is a criterion-free non-parametric method, not compromised by assumptions of data distribution and variance. Its interpretation in binary classifications is similar to accuracy, where a value of 0.5 indicates equal probability of true and false positives and a value of 1 a perfect discrimination between the two conditions. The same cluster-based permutation method implemented for the RSA was used to determine statistical significance (see above).

To determine whether the neural activity patterns that encode face congruency was modulated by Task, classification algorithms were trained and tested to discriminate between congruent and incongruent faces in the sex and emotion task separately. Differences in decoding performance were computed at each time point and participant (AUC emotion – AUC sex/gender). Statistical significance was evaluated using a cluster-based permutation test on these difference scores, following the same procedure described above.

Finally, to test whether expectations based on sex/gender stereotypes improve the encoding of congruent faces compared to incongruent ones, we trained and tested classification algorithms to discriminate between incongruent (happy male vs angry female) and congruent (angry male vs happy female) faces independently. Again, to check for differences (AUC congruent – AUC incongruent), we tested the statistical significance following the same cluster-based permutation test described above.

##### Latency and Slope Analyses

We analyzed differences in the temporal course of decoding (AUC) of congruent vs incongruent faces to evaluate when sex/gender expectations improved performance (see Congruency MVPA section) and the speed of face decoding. To this end, we used complementary latency and slope analyses, to detect when decoding differences emerge, and to estimate differences in the rate of evidence accumulation, i.e., the speed at which neural activity patterns encode faces (Miller et al., 2003; van Ede et al., 2019), respectively. For these analyses, we developed and adapted custom code based on our previously implemented framework (https://github.com/CIMCYC/jacknife-latency-differences).

##### Latency analysis

To determine whether coding of congruent and incongruent faces reaches its maximum at different time points (Miller et al., 2003; van Ede et al., 2019) we compared the latencies of the AUC curves of the first and second peaks, separately. To do this, we smoothed the AUC curves for each condition using a 5-point moving filter, restricted to the 0–1000 ms window. To avoid arbitrarily selecting a single time point (e.g., the peak maximum), we defined nine thresholds relative to the first and second peaks of each curve (from 10% to 90%, in 10% increments). For each of these, we estimated the corresponding latency using a leave-one-out procedure. This was done independently for the first and second peaks and for each condition. For the first peak, the temporal starting point was 0 ms; for the second peak, the intermediate valley between the two peaks was set as the starting point.

We then evaluated latency differences between conditions using a jackknife approach (Miller et al., 2003), which provides a robust estimate of variability across participants. Because this procedure assumes normality, we checked the distribution of latencies at each threshold using the Shapiro–Wilk test (α = 0.05). However, for the second peak, only thresholds between 50% and 90% met this assumption, and thus only these were used. Finally, we applied the Holm–Bonferroni correction to control for multiple comparisons (Miller et al., 2003; van Ede et al., 2019).

##### Slope analysis

This analysis complements previous ones by estimating whether the rate of evidence accumulation differs between the coding of congruent and incongruent faces. A steeper slope indicates faster evidence accumulation, whereas a shallower one reflects slower accumulation.

We calculated slopes by fitting a linear regression model across the different thresholds and their corresponding latencies, independently for each condition and peak. The slope coefficient (β) served as an index of the rate of evidence accumulation towards the maximum decoding value. Then we compared the slopes between conditions using the same jackknife procedure described above.

## 3. Results

### 3.1. Behavioral data

The results (see Figure 2) of the three linear mixed-effect models (see Table 1) were highly consistent across the dependent behavioral variables (MAD, MAD-times EEG RTs). They all showed a significant effect of Congruency (all p < .001) with higher MAD, longer MAD-times and slower RTs during the categorization of incongruent compared to congruent faces. Results of the three analyses were also consistent regarding the effect of task (all ps<.001), with higher MAD, MAD-time and RTs during the emotion compared to the sex task. A significant interaction between Congruency and Task was also found in the three analyses (all ps < .001). Post hoc tests (see Table 2) showed a significant Congruency effect in both tasks, albeit significantly higher in emotion compared to sex judgments (all ps < .001).

**Table 1:** Linear mixed-effects model results for behavioral measures.

| Measure | Predictor | $\beta$ | SE | $df$ | $t$ | $p$ |
| --- | --- | --- | --- | --- | --- | --- |
| **MAD** | Congruency | 52.93 | 5.42 | 61.89 | 9.77 | < .001 |
|  | Task type | 35.55 | 2.42 | 2147.24 | 14.71 | < .001 |
| | Congruency × Task type | 37.00 | 4.83 | 2147.17 | 7.66 | < .001 |
| **MAD-time** | Congruency | 29.32 | 4.22 | 46.62 | 6.95 | < .001 |
|  | Task type | 27.27 | 3.79 | 49.94 | 7.19 | < .001 |
| | Congruency × Task type | 15.06 | 2.76 | 1920.32 | 5.55 | < .001 |
| **RT** | Congruency | 0.042 | 0.008 | 40.23 | 5.19 | < .001 |
|  | Task type | 0.038 | 0.007 | 45.05 | 5.34 | < .001 |
| | Congruency × Task type | 0.023 | 0.004 | 5240.00 | 5.90 | < .001 |

**Note.** $\beta$ = regression coefficient; SE = standard error; $df$ = degrees of freedom; RT = reaction time (ms).

**Table 2:** Post hoc comparisons for the Congruency × Task type interaction across behavioral measures.

| Measure | Contrast | $\beta$ | SE | $z$ | $p$ |
| --- | --- | --- | --- | --- | --- |
| **MAD** | Emotion task: Congruent vs. incongruent | 71.00 | 6.03 | 11.70 | < .001 |
|  | Sex task: Congruent vs. incongruent | 34.20 | 5.83 | 5.86 | < .001 |
| | Contrast ($\beta_{c-e}$ vs. $\beta_{c-s}$) | 36.80 | 4.83 | 7.62 | < .001 |
| **MAD-time** | Emotion task: Congruent vs. incongruent | 36.60 | 4.46 | 8.21 | < .001 |
|  | Sex task: Congruent vs. incongruent | 21.70 | 4.41 | 4.92 | < .001 |
| | Contrast ($\beta_{c-e}$ vs. $\beta_{c-s}$) | 14.90 | 2.76 | 5.41 | < .001 |
| **RT** | Emotion task: Congruent vs. incongruent | 0.054 | 0.008 | 6.44 | < .001 |
|  | Sex task: Congruent vs. incongruent | 0.030 | 0.008 | 3.68 | < .001 |
| | Contrast ($\beta_{c-e}$ vs. $\beta_{c-s}$) | 0.023 | 0.004 | 5.94 | < .001 |

**Note.** Post hoc comparisons were Bonferroni corrected. $\beta_{c-e}$ and $\beta_{c-s}$ refer to congruency effects estimated in the emotion and sex tasks, respectively. RT = reaction time (ms).

### 3.2. RSA

#### 3.2.1. Mouse Trajectories

Full time-resolved model-based RSA (see Figure 3A) revealed significant effects of Congruency (cluster 261 - 792 ms, maximum peak at 522 ms) and Task type (279 - 918 ms, maximum peak around 495 ms). The Emotion and Sex Models also showed significant clusters at early and intermediate stages of processing (Emotion 10 - 558 ms, maximum peak at 413 ms; Sex 17- 737 ms, maximum at 341 ms).

**Figure 3.**
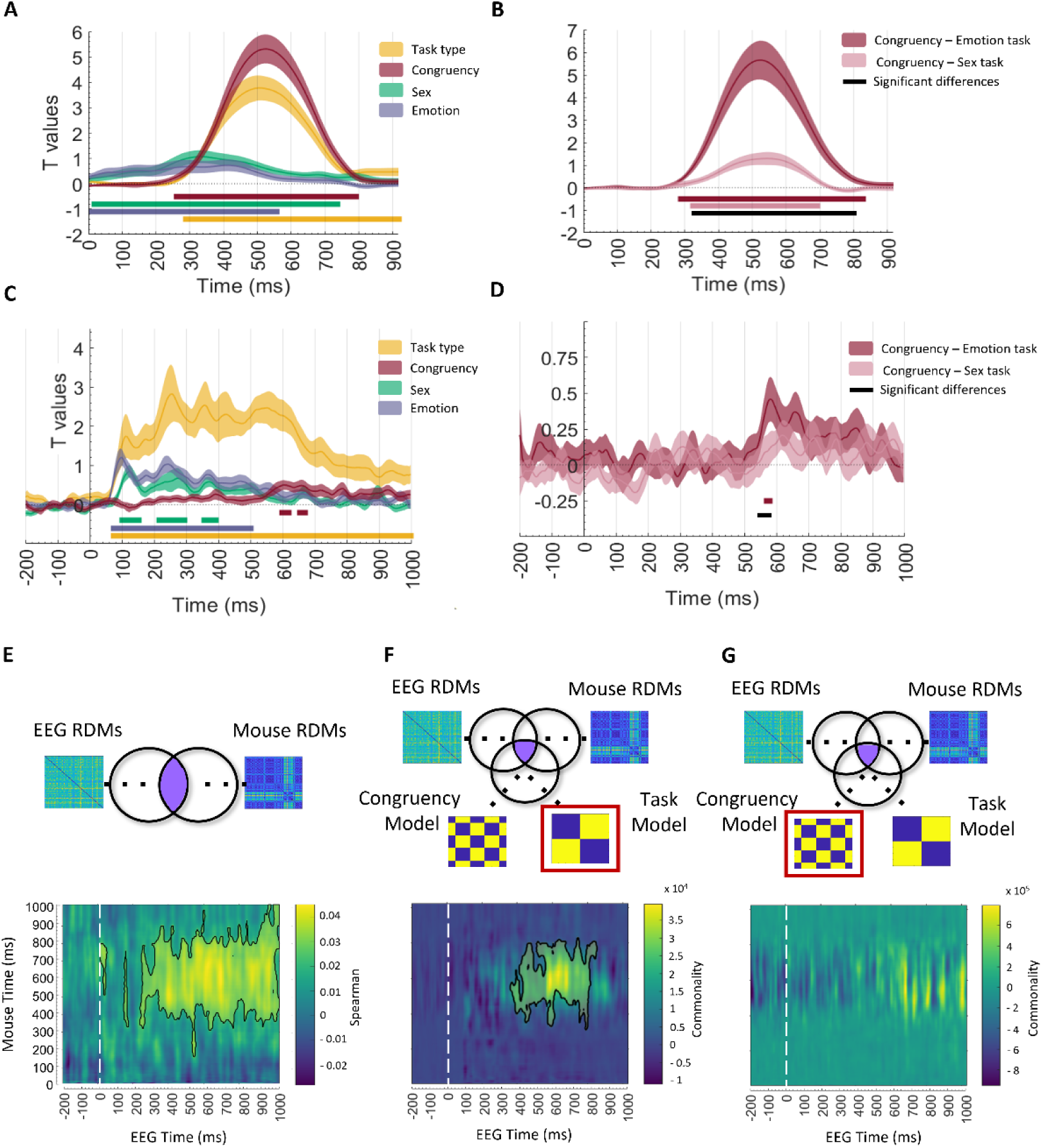
Time-resolved representational similarity analyses (RSA) and fusion EEG–mouse. **(A)** Time-resolved model-based RSA of mouse trajectories using theoretical representational dissimilarity matrices (RDMs) modelling Congruency, Task type, Emotion, and Sex. Curves represent group-level *t*-values across time. Horizontal black bars indicate significant clusters following permutation testing and correction for multiple comparisons. **(B)** Time-resolved RSA of mouse trajectories performed separately for emotion categorization and sex categorization using the Congruency model. The lower panel displays differences between *t*-value curves across task types. **(C)** Time-resolved model-based RSA of EEG activity using theoretical RDMs modelling Congruency, Task type, Emotion, and Sex. Curves represent group-level *t*-values across time, with horizontal black bars indicating significant clusters. **(D)** Time-resolved EEG RSA performed separately for emotion and sex categorization using the Congruency model. The lower panel shows differences between *t*-value curves across task types. **(E)** EEG–mouse simple fusion matrix showing the representational overlap between EEG and mouse trajectories, estimated through Spearman correlations between vectorized RDMs across all combinations of EEG and mouse time points. Color values represent correlation coefficients, and significant clusters are outlined in black. **(F–G)** EEG-Mouse fusion model-based commonality analyses showing the variance shared between EEG and mouse representational geometries explained by Task type (**F**) and Congruency (**G**). Color scales represent commonality coefficients, and black contours indicate significant clusters.

The results of the RSAs separately for each Task (see Figure 3B) revealed a significant effect of Congruency at intermediate and late stages of the trajectories, in both emotion (280 - 840 ms, maximum peak at 530 ms) and sex (325 - 700 ms, maximum peak at 550 ms) tasks. A comparison between their T-values curves showed a significantly higher Congruency effect in the emotion task at intermediate and late stages (320 – 810 ms).

#### 3.2.2. EEG data

Time-resolved model-based RSA of EEG data (see Figure 3C) revealed a significant effect of Congruency (two continuous clusters between 597 and 668 ms, with a maximum peak at around 656 ms). The Task model showed a significant effect throughout the epoch (from 74 ms until the end of the epoch, with a maximum peak around 254 ms). The results showed a significant early and intermediate effect for Emotion and Sex models (Emotion 74 - 503 ms, maximum peak at 103 ms; Sex 100 - 390 ms, maximum at 128 ms).

The results of the RSA on the emotion task showed a significant effect of Congruency model at intermediate stages of the epoch (570 - 580 ms, with a maximum peak at around 574 ms; see Figure 3 D). However, Congruency was not significant in the RSA during the sex categorization task. Finally, the difference between T-values curves showed a significantly higher impact of Congruency in the emotion task (570 – 588 ms).

### 3.3. EEG-Mouse Fusion

To evaluate the overlap in representational structure between the spatiotemporal patterns of mouse trajectories and EEG data, we conducted a simple mouse-EEG fusion based on Spearmańs correlations. As shown in Figure 3E, the results revealed similarity with significant clusters spanning approximately 400–900 ms in mouse trajectory time and 250–1000 ms in EEG time.

Model-based fusion results revealed that the shared variance was explained by the Task model (see Figure 3F; approximately 400 - 800 ms in mouse trajectories and 350 - 800 ms in EEG). In contrast, we found no evidence for the Congruence model (see Figure 3G).

### 3.4. Time resolved Multivariate Classifiers

Time-resolved multivariate classifiers tested whether and when neural coding was modulated by expectations and task context, and whether such coding is more robust for faces congruent with sex/gender stereotypes.

Congruency was encoded progressively along the trial epoch, with a maximum peak around 500 ms (see Figure 4A). Coding of task context showed a fast and sustained increase and peaks at different stages (117, 281 and 457 ms). Facial emotion and sex also impacted neural coding during early and intermediate processing windows, with a first maximum peak around 90 – 110 ms (see Figure 4A).

**Figure 4.**
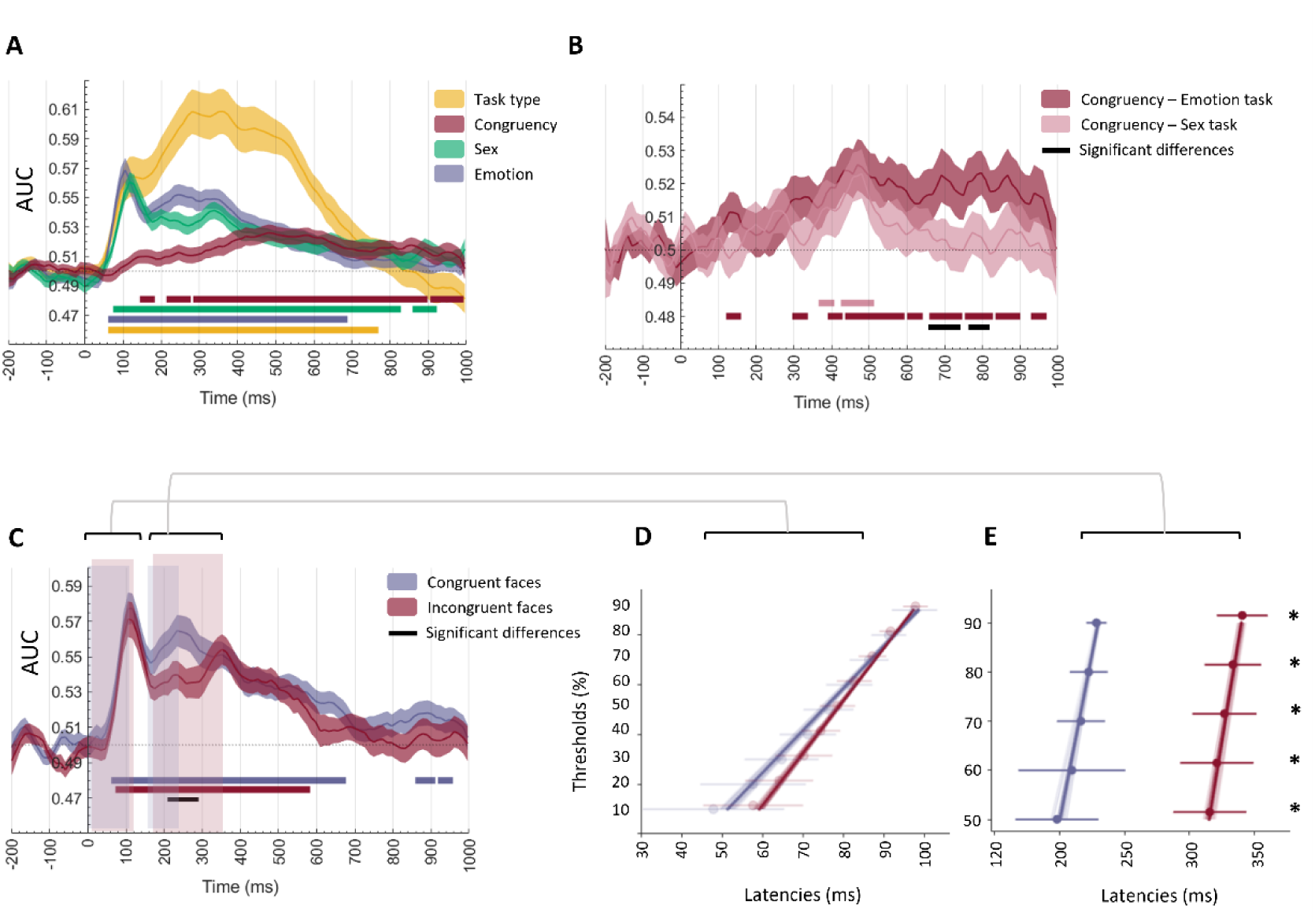
Time-resolved Multivariate Classifiers of face congruency, task context, and latencies-slopes analysis. **(A)** Time-resolved multivariate pattern classification (MVPA) of EEG activity decoding Congruency, Task type, Emotion, and Sex. Curves represent group-level decoding performance quantified as area under the curve (AUC) across time. Horizontal colors bars indicate significant above-chance decoding following permutation testing and correction for multiple comparisons. **(B)** Time-resolved decoding of face Congruency performed separately for emotion categorization and sex categorization tasks. Horizontal colors bars indicate significant above-chance decoding following permutation testing and correction for multiple comparisons. The lower panel displays differences in decoding performance (*AUC* emotion − *AUC* sex) across time, with horizontal black bars indicating significant clusters. **(C)** Time-resolved decoding of congruent faces (angry male vs. happy female) and incongruent faces (happy male vs. angry female). Curves represent group-level AUC values across time. Horizontal colors bars indicate significant above-chance decoding following permutation testing and correction for multiple comparisons. The lower panel shows differences in decoding performance between congruent and incongruent faces, with significant clusters indicated by horizontal black bars. **(D)** Latency and slope analyses derived from congruent and incongruent decoding curves. Latency analyses quantified the temporal emergence of neural decoding across thresholds relative to peak AUC values, whereas slope analyses estimated the rate of evidence accumulation toward peak decoding. Error bars represent confidence intervals, and asterisks indicate statistically significant differences following correction for multiple comparisons.

Classifiers applied separately to each task showed progressive neural coding of Congruency, which was consistent until late stages during the emotion discrimination but showed a progressive decline toward the end of the epoch during the sex task. The difference between AUC curves revealed significantly higher Congruency decoding in emotion in mid to late stages of the epoch (668-797 ms; see Figure 4B).

On the other hand, neural activity patterns encoded congruent (angry male vs. happy female) and incongruent (happy male vs angry female) faces through early, middle and late stages of processing. Whereas both AUC value curves showed a first maximum peak at approximately 110 ms, a second maximum peaked earlier for congruent (250 ms) than incongruent faces (351 ms). Comparisons between the AUC curves revealed higher coding accuracy for congruent faces from 234 to 281 ms (see Figure 3C).

### 3.5. Latencies and Slopes

To evaluate whether expectations improve the efficiency and speed with which neural activity patterns encode facial stimuli, we applied a jackknife procedure to the latencies and slopes of the AUC curves from the previous multivariate classifiers (congruent and incongruent faces). The results of latencies and slopes to the first maximum peak did not reveal significant differences between congruent and incongruent AUC curves at any thresholds (all p > 0.05). In contrast, the analyses to the second maximum peak (see Table 4 and Figure 4E) showed significantly earlier latencies for congruent (M_avg_ = 214.88 ms) than incongruent faces (M_avg_ = 327.64 ms) at all thresholds (all p < 0.001). There were no differences between the slopes (all ps > 0.05; see Table X and Figure 4D).

**Table 3:**
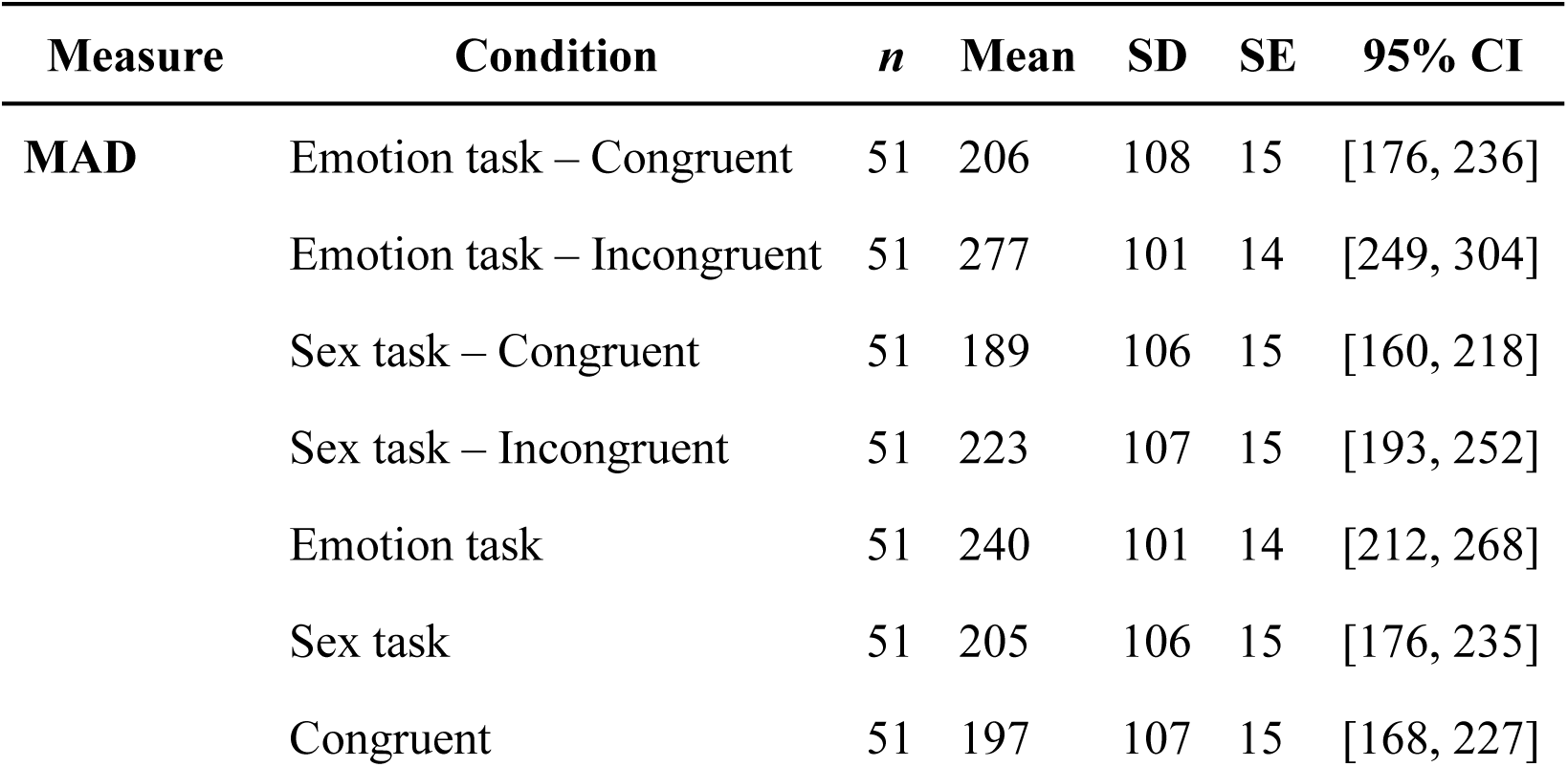
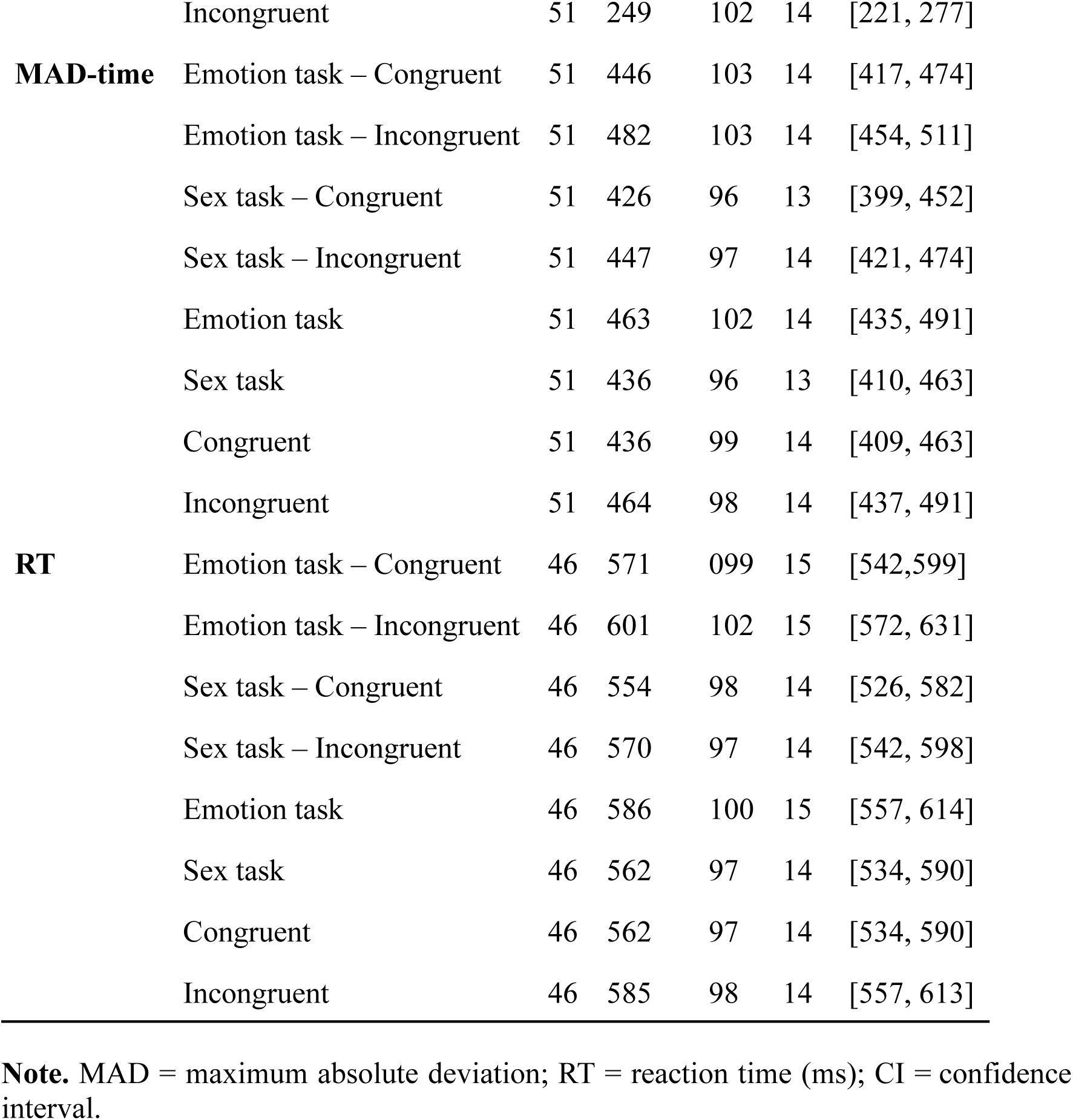
Descriptive statistics for behavioral measures across tasks and congruency conditions.

**Table 4:** Latency differences between congruent and incongruent face decoding across thresholds.

| Peak | Threshold | Mean latency congruent (ms) | CI congruent (ms) | Mean latency incongruent (ms) | CI incongruent (ms) | Difference (ms) | CI difference (ms) | *p* | Corrected *p* |
| --- | --- | --- | --- | --- | --- | --- | --- | --- | --- |
| 1 | 0.1 | 47.87 | [30.44, 65.30] | 57.66 | [45.35, 69.98] | −9.79 | [−32.26, 12.68] | .384 | 1.000 |
|  | 0.2 | 57.66 | [44.63, 70.68] | 64.07 | [55.68, 72.46] | −6.41 | [−23.10, 10.27] | .442 | 1.000 |
|  | 0.3 | 64.69 | [55.56, 73.82] | 70.14 | [63.08, 77.21] | −5.45 | [−17.93, 7.02] | .383 | 1.000 |
|  | 0.4 | 71.14 | [64.19, 78.09] | 74.34 | [69.52, 79.17] | −3.21 | [−12.06, 5.65] | .469 | 1.000 |
|  | 0.5 | 76.26 | [70.10, 82.43] | 78.45 | [74.08, 82.81] | −2.19 | [−9.86, 5.49] | .568 | 1.000 |
|  | 0.6 | 81.38 | [75.66, 87.10] | 82.60 | [78.30, 86.89] | −1.22 | [−8.27, 5.83] | .728 | 1.000 |
|  | 0.7 | 86.27 | [83.67, 90.48] | 87.08 | [83.67, 90.48] | −0.81 | [−6.54, 4.92] | .777 | 1.000 |
|  | 0.8 | 91.12 | [86.84, 95.41] | 91.55 | [88.80, 94.31] | −0.43 | [−5.40, 4.54] | .862 | 1.000 |
|  | 0.9 | 97.48 | [94.66, 100.74] | 97.70 | [94.66, 100.74] | −0.22 | [−6.52, 6.09] | .945 | 1.000 |
| 2 | 0.5 | 197.89 | [165.85, 229.93] | 315.73 | [287.53, 343.92] | −117.84 | [−162.27, −73.41] | < .001 | < .001 |
|  | 0.6 | 209.37 | [168.13, 250.61] | 321.19 | [292.95, 349.44] | −111.82 | [−164.59, −59.05] | < .001 | < .001 |
|  | 0.7 | 216.41 | [191.72, 241.10] | 327.01 | [302.30, 351.72] | −110.59 | [−146.00, −75.19] | < .001 | < .001 |
|  | 0.8 | 222.39 | [207.77, 237.02] | 333.54 | [311.70, 355.37] | −111.14 | [−141.72, −80.57] | < .001 | < .001 |
|  | 0.9 | 228.37 | [220.58, 236.10] | 340.73 | [321.10, 360.35] | −112.36 | [−135.83, −88.89] | < .001 | < .001 |

**Note.** Latencies were estimated relative to thresholds defined as proportions of peak decoding performance (AUC). Corrected *p*-values correspond to Holm–Bonferroni adjustments.

## 4. Discussion

A common tenet in predictive processing is that expectations facilitate the encoding of expected information. Our findings suggest that these facilitatory effects are not uniform during face perception; rather, their timing and magnitude are flexibly modulated by task goals. More broadly, these results indicate that the influence of prior knowledge on perception is dynamically adjusted according to current behavioral requirements, supporting the view that predictive processes serve an adaptive function beyond the mere facilitation of expected information. Using a combined mouse-tracking and EEG approach, we found that expectations derived from sex/gender stereotypes modulate behavioral performance and neural representational dynamics during face processing. Importantly, these effects were significantly stronger during emotion categorization than during sex judgments, indicating that perceptual expectations exert greater influence when processing relies more heavily on top-down computations. In addition, the facilitatory effect associated with congruent faces was reflected not only in enhanced neural decoding, but also in earlier decoding latencies. This suggests that expectations facilitate face processing not by increasing the rate of perceptual evidence accumulation, but by accelerating the temporal emergence of neural coding patterns.

Our results consistently show that expectations facilitate processing of faces. At the behavioral level, incongruent faces elicited larger MAD, longer MAD-times and slower RTs relative to congruent ones. These results replicate previous mouse tracking studies (Barnett et al., 2021; Gutiérrez-Blanco et al., 2025; Stolier & Freeman, 2016) and align with behavioral findings on expectation-based facilitation (Becker et al., 2007; Falbén et al., 2022; Primbs et al., 2022). Converging evidence emerged from multivariate neural analyses. RSA revealed differences between the spatiotemporal patterns of mouse trajectories and the neural representational space of EEG data when processing incongruent versus congruent faces. In line with these, neural activity patterns reflect congruency from early perceptual stages starting at approximately ∼150 ms, as shown by classifiers’ decoding performance increasing progressively over time. Furthermore, neural patterns were more reliable for congruent than incongruent faces between 200 and 300 ms. Our findings extend previous predictive processing neuroimaging studies (Barnett et al., 2021) into the temporal domain, suggesting that the top-down modulation exerted by higher-order regions onto face-sensitive perceptual areas unfolds from early stages of face processing.

Although previous findings show that expectations facilitate early face processing, they do not clarify the underlying temporal mechanisms. Our latency analyses revealed that congruent faces reached peak neural discriminability approximately 110 ms earlier than incongruent faces. In contrast, we found no differences in the slopes of their decoding curves, indicating that the rate of perceptual evidence accumulation remained comparable across conditions. Together, these findings suggest that expectations based on sex/gender stereotypes facilitate face processing by accelerating the temporal emergence of neural coding patterns rather than by increasing the speed of evidence accumulation. This interpretation is consistent with predictive processing accounts proposing that prior expectations preactivate sensory templates of expected information through top-down signaling (Kok et al., 2017), modulating perceptual systems toward neural states partially aligned with expected stimuli. To our knowledge, this is the first study to directly dissociate these two temporal mechanisms during predictive face processing, distinguishing between accelerated emergence of neural coding patterns and changes in evidence accumulation dynamics. Beyond extending previous work on social expectations and face perception, this approach may provide a useful framework for future studies investigating how perceptual predictions shape the temporal dynamics of neural information processing.

A central aspect of our study was to determine whether different task demands, related to the processing of variant (emotion expression) and invariant (sex) facial features could modulate the facilitatory effect of expectations. Our results were consistent with this. Across behavioral and neural analyses, expectation effects were higher for emotion than for sex categorization. Behaviorally, the impact of incongruent faces was consistently larger during emotion categorization, reflected in slower reaction times, larger trajectory deviations (MAD) and longer MAD-times relative to sex categorization. This same pattern emerged at the neural level, where both RSA and classifiers revealed stronger congruency effects during emotion than sex judgements. This convergence suggests that emotional expression processing is more strongly influenced by expectations than facial sex processing. These findings are consistent with previous evidence showing an asymmetric relationship between emotion and sex categorization (Atkinson et al., 2005; Karnadewi & Lipp, 2011), with emotion depending more strongly on top-down modulation and sex categorization on bottom-up perceptual input (Gandolfo et al., 2025). Together, our findings suggest that perceptual expectations do not uniformly apply during face processing but are rather flexibly weighted according to the computational demands of different categorization goals. Specifically, the stronger dependence of emotional expression processing on top-down modulation may render it especially susceptible to the influence of expectations derived from sex/gender stereotypes. Our findings provide novel evidence for predictive processing accounts by suggesting that the impact of perceptual expectations depends on the computational architecture of the perceptual process itself, exerting stronger effects on perceptual operations that rely more heavily on top-down signaling.

A further objective of the present study was to determine the similarity between the representational geometric structures of mouse trajectories and EEG data during face perception, and the part of this potential overlap that is explained by task demands and by expectations. Simple EEG-mouse fusion analyses revealed significant similarity between the geometries of both modalities during intermediate and late stages of processing, suggesting that during face processing, both representational structures reflect similar information processing. Findings are consistent with previous work showing similarities between mouse trajectories and MEG data during object and face perception (Koenig-Robert et al., 2024) and extend the application of these multimodal representational approaches to the field of predictive processing. Compared to previous fMRI studies (Barnett et al., 2021; Stolier & Freeman, 2017), this approach provides a finer characterization of the temporal dynamics through which expectations unfold across neural and behavioral domains. Importantly, model-based fusion analyses revealed that the shared variance between EEG and mouse data was primarily explained by task demands, whereas no significant shared variance emerged for face congruency. This pattern does not indicate that congruency effects were absent in either modality, rather it suggests that the cognitive mechanisms underlying each modality reflect congruency differently. One possibility is that the neural representational space of EEG data reflects perceptual modulations of congruency, whereas the mouse spatiotemporal patterns of trajectories reflect processes more closely linked to decision-making and motor responses. More broadly, these findings highlight the potential of multimodal representational approaches.

Despite the novel findings of the present study, several limitations should be acknowledged. As the experimental design did not include a neutral control condition, the present results do not allow determining whether congruency primarily reflects facilitation or interference, or a combination of both. Future studies incorporating neutral emotional expressions or androgynous faces may help disentangle these possibilities. A second limitation concerns the lack of spatial precision inherent to EEG measures, so the present findings do not allow direct identification of the neural regions and pathways underlying the dynamics uncovered. Future multimodal neuroimaging studies combining high temporal and spatial resolution techniques could determine whether the stronger expectation effects observed during emotion categorization are preferentially associated with dorsal face-processing systems and greater top-down connectivity.

## 5. Conclusions

Do expectations influence neural coding in the same way regardless of the task at hand? Our results suggest they do not. The present study provides converging behavioral and neural evidence that expectations derived from sex/gender stereotypes facilitate face processing flexibly, according to task demands, with stronger influence during coding of emotional information. In addition, perceptual expectations facilitate face processing by accelerating the temporal emergence of neural coding patterns rather than by increasing the rate of evidence accumulation. In summary, our findings support a model in which predictions are dynamically adjusted to the goals of the task at hand.

## Author Contributions

Author contributions: F.G-B., A.F.P., C.G.-G., and M.R. designed research; F.G-B and S.G recall data. F.G-B performed research; F.G-B., A.F.P., C.G.-G., and M.R. contributed unpublished reagents/analytic tools; F.G-B. analyzed data; F.G-B., A.F.P., C.G.-G. and M.R. wrote the paper.

## Ethics statement

The authors declare that participants provided their informed written consent to participate in this study, prior to the experiment, which took place at the Mind, Brain and Behavior Research Centre (CIMCYC). The study was approved by the Ethics Committee of the University of Granada (1584/CEIH/2020).

## Data Availability

The data and analysis code supporting the findings of this study will be made publicly available upon publication.

## Acknowledgements

We are grateful to Sergio Gaspar for his assistance with data collection and Jose.M.G. Peñalver for his assistance with the EEG equipment setup.

